# Estrogen signaling in the dorsal raphe regulates binge-like drinking in mice

**DOI:** 10.1101/2022.09.05.506646

**Authors:** Valeria C. Torres Irizarry, Bing Feng, Xiaohua Yang, Patel Nirali, Sarah Schaul, Lucas Ibrahimi, Hui Ye, Pei Luo, Leslie Carrillo-Sáenz, Penghua Lai, Maya Kota, Devin Dixit, Chunmei Wang, Amy W. Lasek, Yanlin He, Pingwen Xu

## Abstract

The ovarian hormone estrogens promote binge alcohol drinking and contribute to sex differences in alcohol use disorder. However, the mechanisms for estrogen-induced binge drinking are largely unknown. This study aims to test if estrogens act on 5-hydroxytryptamine neurons in the dorsal raphe nucleus (5-HT^DRN^) to promote binge drinking. We used the drinking in the dark (DID) behavioral test in mice to mimic binge drinking in humans. We found that female mice drank more alcohol than male mice in chronic DID tests. This sex difference was associated with distinct alterations in mRNA expression of estrogen receptor α (ERα) and 5-HT-related genes in the DRN, suggesting a potential role of estrogen/ERs/5-HT signaling in binge alcohol drinking. In supporting this view, 5-HT^DRN^ neurons from naïve male mice had lower baseline neuronal firing activity but higher sensitivity to alcohol-induced excitation compared to 5-HT^DRN^ neurons from naïve female mice. Notably, this higher sensitivity was blunted by 17β-estradiol treatment in males, indicating an estrogen-dependent mechanism. We further showed that both ERα and ERβ are expressed in 5-HT^DRN^ neurons, whereas ERα agonist propyl pyrazole triol (PPT) depolarizes 5-HT^DRN^ neurons and ERβ agonist diarylpropionitrile (DPN) hyperpolarizes 5-HT^DRN^ neurons. Notably, both PPT and DPN treatments blocked the stimulatory effects of alcohol on 5-HT^DRN^ neurons in males, despite the fact that they have antagonistic effects on the activity dynamics of 5-HT^DRN^ neurons. These results suggest that ERs’ inhibitory effects on ethanol-induced burst firing of 5-HT^DRN^ neurons may contribute to higher levels of binge drinking in females. Consistently, chemogenetic activation of ERα- or ERβ-expressing neurons in the DRN reduced binge alcohol drinking. These results support a model in which estrogens act on ERα/β to prevent alcohol-induced activation of 5-HT^DRN^ neurons, which in return leads to higher binge alcohol drinking.

## Introduction

Binge drinking, defined as the consumption of a significant amount of alcohol in a single setting^1,2^, is the most common pattern of excessive alcohol use in the US^3^ and contributes to the development of alcohol use disorder (AUD). Like many other psychiatric disorders, there are sex differences in the trajectory of AUD and its consequences. Binge drinking by women has increased faster than by men in recent decades^4^ and women show more vulnerability to alcohol-induced cognitive impairment and peripheral neuropathy than men^5-9^. While more effective treatments to reduce alcohol misuse in both sexes are urgently needed, our understanding of the pathophysiology of female binge drinking is limited.

A large body of evidence indicates that the ovarian hormone estrogens play a role in sex differences in alcohol consumption behavior. Human clinical studies support the association of increased estrogen levels and increased alcohol use^10-13^. Consistently, numerous animal studies have demonstrated the stimulatory effects of 17β-estradiol (E2) on ethanol drinking under various access conditions, including those that promote binge drinking^14-19^. Overall, human and animal studies implicate estrogens in increased alcohol drinking. However, little is known about the central mechanisms mediating estrogenic regulation of behaviors related to alcohol drinking. Only a handful of studies have shown that ERs in the ventral tegmental area mediate estrogen’s effects on the dopamine system and increase alcohol binge-like drinking behavior in female mice^20-22^. Nevertheless, the other potential ERs sites in the brain mediating estrogenic effects on alcohol binge drinking have not been identified.

The brain serotonin (5-hydroxytryptamine, 5-HT) system is a critical modular of alcohol intake and critically involved in alcohol’s effects on the brain and the development of alcohol misuse ^23-25^. It has been reported that binge drinking induces a burst release of central 5-HT, and increased brain 5-HT content inhibits alcohol consumption in both humans and rodents^23-26^. Interestingly, estrogens have a potent modulatory impact on this system. For example, estrogens have been shown to increase serotonergic tone by regulating the synthesis and degradation of serotonin^27-29^. In line with these findings, both estrogen receptor α (ERα) and β (ERβ) are highly expressed in the dorsal raphe (DRN)^30^, the largest serotonergic nucleus and a major source of 5-HT. It has been shown that the ERα agonist propyl pyrazole triol (PPT) increases^31,32^ whereas the ERβ agonist diarylpropionitrile (DPN) decreases the spontaneous firing of the serotonergic neurons in the DRN^33^. These studies demonstrate a potential mediating role of DRN 5-HT neurons (5-HT^DRN^) in the regulatory effects of estrogen on binge drinking.

In this study, we first used a chronic drinking in the dark (DID) behavioral test to examine the effects of long-term binge drinking on estrogen and 5-HT signaling in the DRN. Furthermore, the *ex vivo* responsiveness of 5-HT^DRN^ neurons to ethanol treatment was compared between male and female mice using whole-cell patch-clamp electrophysiology recording. We further tested whether E2, PPT, and DPN treatments attenuate the sex difference in ethanol-induced activity changes of 5-HT^DRN^ neurons. Finally, we used the designer receptors exclusively activated by designer drugs (DREADD) approach to examine the effects of DRN neural activity on binge drinking in mice. Our results provide compelling evidence to support a model in which estrogens act on ERα/β to prevent alcohol-induced activation of 5-HT^DRN^ neurons that inhibit binge drinking.

## Results

### Sex-specific gene changes in the DRN induced by chronic DID

Both male and female C57BL/6J mice were subjected to 9-week water or ethanol DID. We found females consumed more alcohol than males in the ethanol, but not water DID test (Fig. 1A-D). Notably, 9-week binge-like ethanol drinking leads to increases in the mRNA expression of estrogen receptor 1 (*Esr1*, the gene encoding ERα protein), serotonin transporter (*Sert*, serotonin reuptake enzyme), and plasmacytoma expressed transcript 1 (*Pet1*, a crucial transcription factor for the metabolism and the reuptake of serotonin) in males. Conversely, chronic binge-like drinking decreased the mRNA expression of *Esr1* but not 5-HT-related genes in females (Fig. 1E-J), suggesting a sexually dimorphic response in the DRN to chronic binge-like ethanol drinking.

**Fig. 1.**
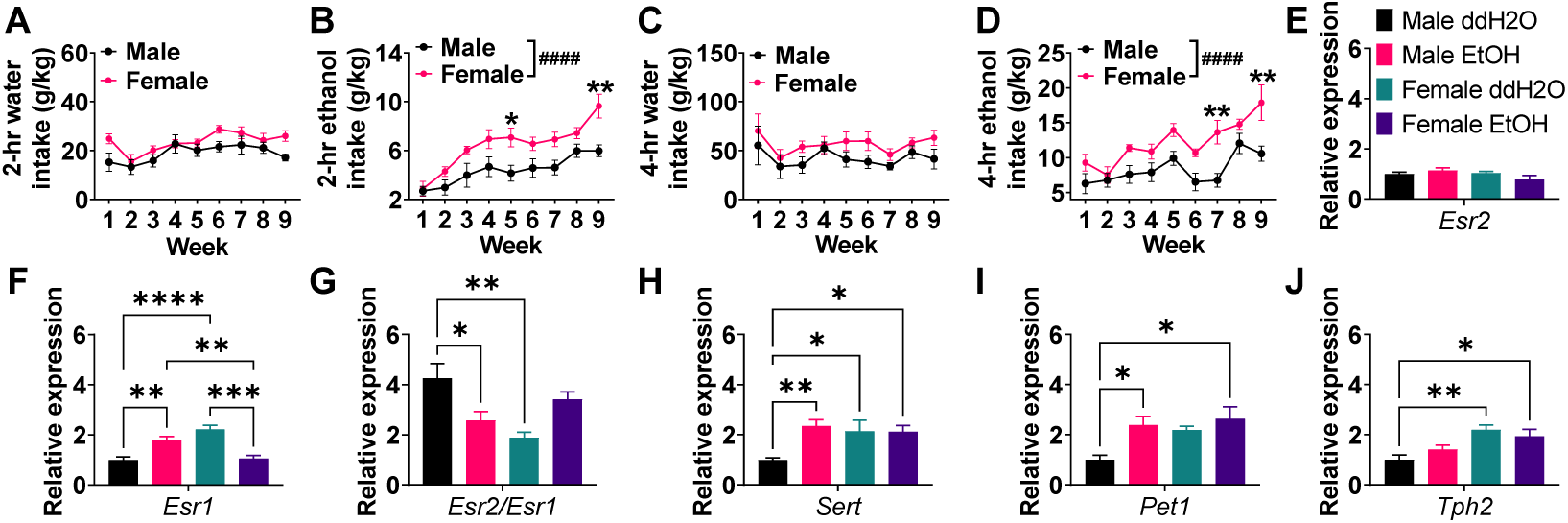
Chronic binge-like ethanol drinking leads to sex-specific alterations in Esr1 and 5-HT-related genes in the DRN of mice. (A-B) Average water (A) or EtOH (B) consumed during 2-hr drinking sessions on days 1 to 4 each week (n = 5 or 7). (C-D) Average water (C) or EtOH (D) consumed during 4-hr drinking sessions on day 4 each week (n = 5 or 7). (E-J) Relative mRNA expression of estrogen receptor 2 (Esr2, E), estrogen receptor 1 (Esr1, F), Esr1/Esr2 (G), serotonin transporter (Sert, H), plasmacytoma expressed transcript 1 (Pet1, I), and tryptophan hydroxylase 2 (Tph2, J) in the DRN (n = 4-6). Mean ± SEM. (A-D) #### p<0.0001 in two-way ANOVA analysis, *p<0.05, **p<0.01 in the following post hoc Sidak tests. (E-J) *p<0.05, **p<0.01, ***p<0.001, ****p<0.0001 in one-way ANOVA analysis followed by post hoc Tukey tests.

### 5-HT^DRN^ neurons co-express ERα/ERβ

To examine the expression pattern of ERα and ERβ on serotonergic neurons in the DRN, we did dual immunofluorescent (IF) staining of ERα and tryptophan hydroxylase (TPH) in the *Esr2*-Cre/Rosa26-LSL-tdTOMATO mice, in which all ERβ positive neurons were labeled with red TOMATO fluorescence. We found that about 80-90% of 5-HT^DRN^ neurons co-express ERα or ERβ (83.63% in M vs. 89.33% in F); about 40-50% of 5-HT^DRN^ neurons are positive for both ERα and ERβ (41.61% in M vs. 53% in F, Fig. 2A-C). Notably, there are more ERα (+) 5-HT^DRN^ neurons than ERβ (+) 5-HT^DRN^ neurons in both males and females (77.4% vs. 47.84% in M; 79.2% vs. 63.13% in F, Fig. 2C). We also found that around 80% of ERα^DRN^ neurons (85.28% in M vs. 79.48% in F) and 70% of ERβ^DRN^ neurons (70.26% in M vs. 73.02% in F; Fig. 2D-E) are serotonergic. Consistently, we found abundant ERα and ERβ expression in 5-HT neurons in the DRN of ERβ-EGFP mice, as indicated by white arrow-pointed triple color-positive neurons (Fig. 2F-G). These provide the anatomic basis that estrogen acts through both ERα/β expressed by 5-HT^DRN^ neurons to regulate binge drinking behavior.

**Fig. 2.**
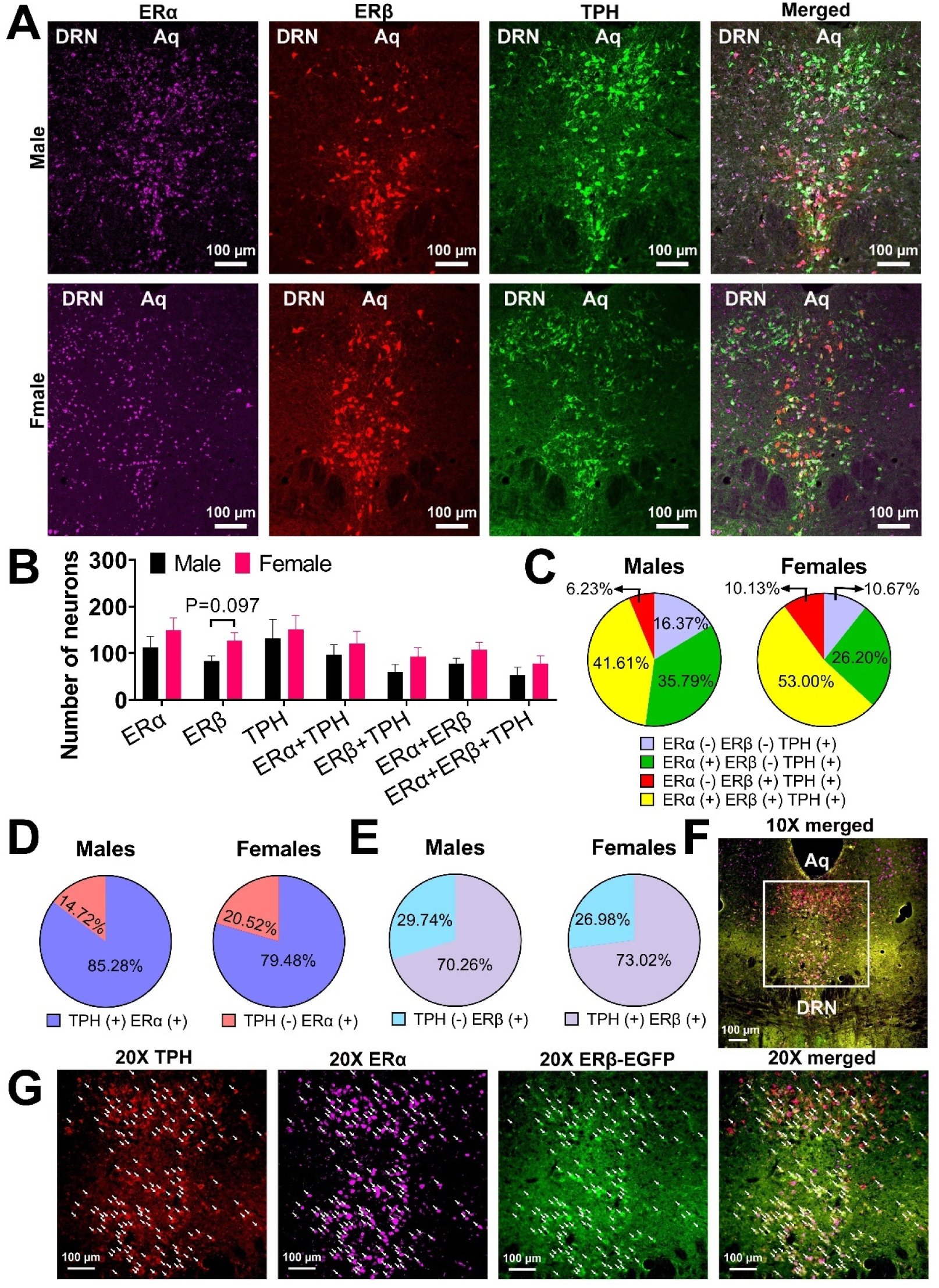
ERαβ are expressed in 5-HT^DRN^ neurons. (A) Double immunofluorescent staining for ERα (purple) and TPH (green) in the DRN of *Esr2*-Cre/Rosa26-tdTOMATO mice. (B) Summary of quantification per section (n = 3 and 3). (C) Percentage of ERα and ERβ positivity in TPH^DRN^ neurons. (D-E) Percentage of TPH positivity in ERα^DRN^ (D) or ERβ^DRN^ (E) neurons. (F-G) Low (F) and high magnification of TPH (red, G), ERα (purple), ERβ-EGFP (green), and merged (triple) in the DRN of a female ERβ–EGFP mouse. White arrows point to triple positive TPH (+) ERα (+) ERβ (+) neurons. Mean ± SEM.

### Sexually dimorphic responses to ethanol in ERα^DRN^ neurons

To determine the baseline characteristics of ERα^DRN^ neurons, we recorded DRN ZsGreen(+) neurons in *ex vivo* brain slices from ERα-ZsGreen mice (Fig. 3A). We found that ERα^DRN^ neurons from female mice showed higher firing frequency and depolarized resting membrane potential compared to ERα^DRN^ neurons from male mice (Fig. 3B-D), indicating a higher spontaneous neural activity in ERα^DRN^ neurons. We further tested the dose responses of ERα^DRN^ neurons to ethanol treatment in both sexes. ERα^DRN^ neurons from males were more sensitive to ethanol-induced excitation compared to ERα^DRN^ neurons from females (Fig. 3E-J). Specifically, while a low dose of 0.5 mM ethanol treatment failed to increase the firing frequency of female ERα^DRN^ neurons (Fig. 3F), it significantly raised the firing rate of male ERα^DRN^ neurons (Fig. 3E). Consistently, the ethanol-induced increases in firing frequency were substantially more significant in males than in females at 0.5 and 50 mM doses (Fig. 3G). The changes in resting membrane potential were also greater in males at a dose of 1 mM ethanol treatment compared to female mice (Fig. 3J). Since a majority of ERα-expressing cells within the DRN are 5-HT neurons (Fig. 2D)^32^, we speculate that ERα-expressing 5-HT^DRN^ neurons exhibit a sexually dimorphic response to ethanol treatment.

**Fig. 3.**
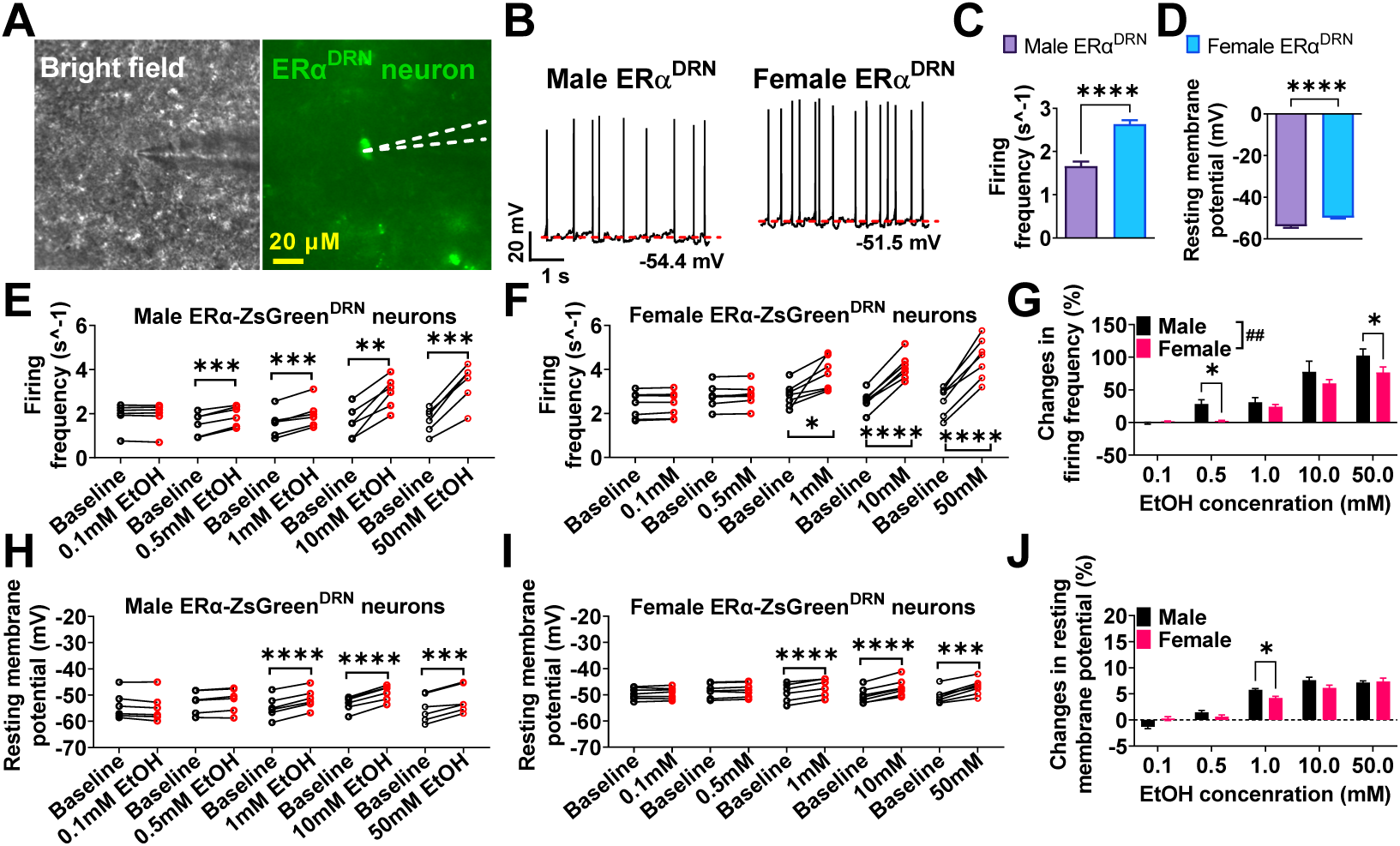
Sex difference in the responses of ERα^DRN^ neurons to ethanol. (A) Micrographic images showing a recorded ERαZsGreen (+) neurons in the DRN of female mice. (B-D) Representative electrophysiological trace (B), baseline firing frequency (C), and resting membrane potential (D) of ERα-ZsGreen (+) neurons in the DRN of male and female ERα-ZsGreen mice (n = 30 or 35). (E-J) The dose-dependent responses of firing frequency and resting membrane potential (n = 6 or 7) before and after a 1s puff of aCSF containing different doses of EtOH. Mean ± SEM. (C-D) ****p<0.0001 in unpaired t-tests. (E-F and H-I) *p<0.05, **p<0.01, ***p<0.001, ****p<0.0001 in paired t-tests. (G and J) ##p<0.01 in two-way ANOVA analysis, *p<0.05 in the following post hoc Sidak tests.

### Estrogen attenuates ethanol-induced excitation of 5-HT^DRN^ neurons in males

To compare the baseline characteristics of 5-HT^DRN^ neurons from different sexes, we recorded DRN tdTOMATO(+) neurons in *ex vivo* brain slices from TPH2-iCreER/Rosa26-LSL-tdTOMATO mice (Fig. 4A). Similar to what we observed in ERα^DRN^ neurons, female 5-HT^DRN^ neurons had a higher spontaneous activity, as indicated by increased firing frequency and depolarized resting membrane potential (Fig. 4B-D). To further test if estrogens contribute to sex differences in ethanol’s regulation of DRN neuron firing, we recorded the responses of 5-HT^DRN^ neurons to 1s puff treatment of 1 mM ethanol in the presence of vehicle or E2. Consistent with ethanol’s dose-dependent stimulatory effects on ERα^DRN^ neurons (Fig. 3E-J), 1 mM ethanol puff significantly increased firing frequency and depolarized resting membrane potential of 5-HT^DRN^ neurons regardless of sex or E2 treatment (Fig. 4E-F and I-J). Notably, at a baseline level without ethanol treatment, E2 incubation did not affect the firing rate or resting membrane potential of 5-HT^DRN^ neurons in both sexes (Fig. 4G and K). However, the ethanol-induced increases in firing frequency and resting membrane potentials were significantly reduced in males but not in females by E2 treatment (Fig. 4H and L), suggesting an attenuation in ethanol-induced activation of 5-HT^DRN^ neurons.

**Fig. 4.**
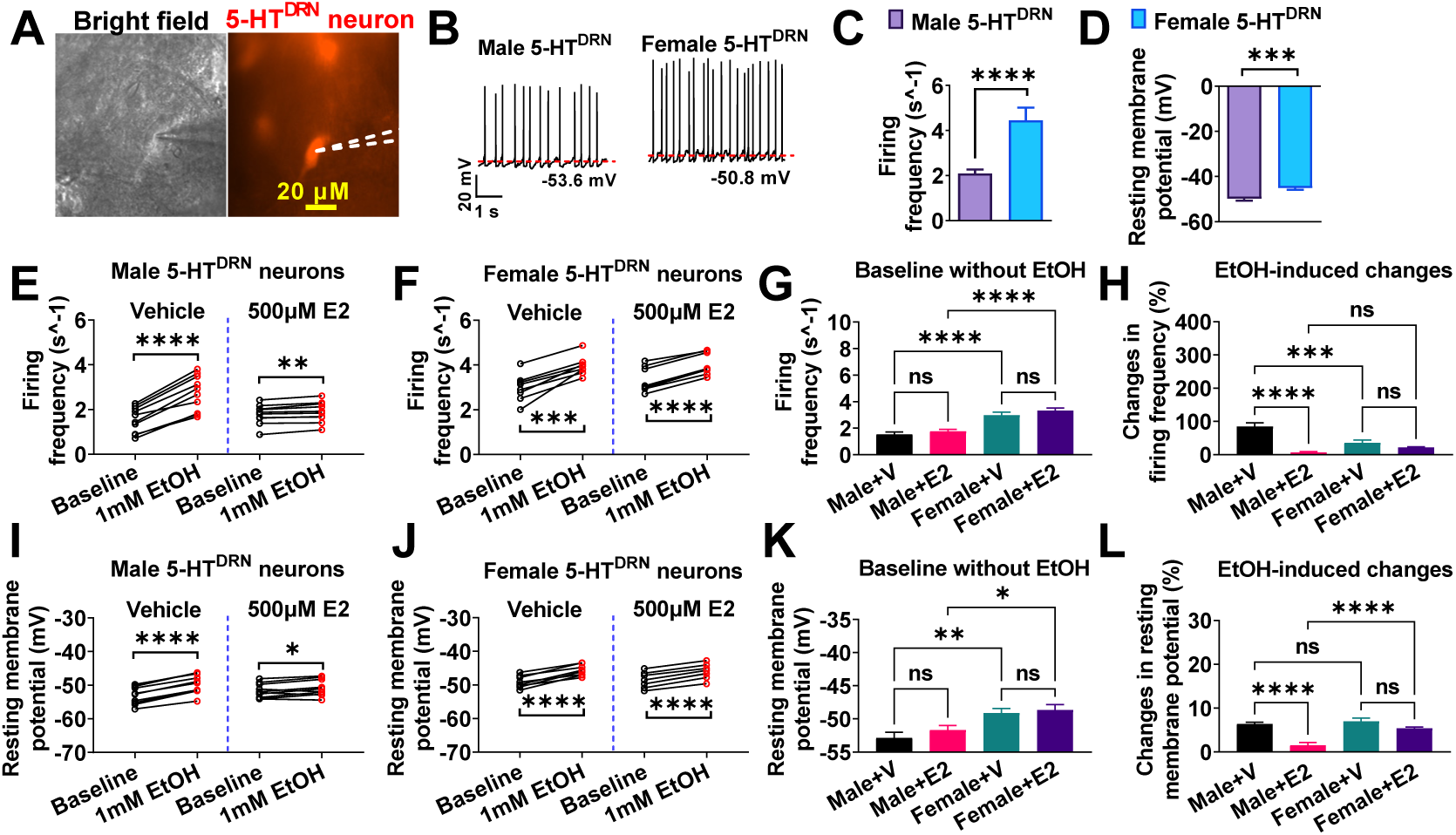
Estrogen attenuates EtOH-induced activation of 5-HT^DRN^ neurons. (A)Micrographic images a recorded TPH2 (+) neuron in the DRN of female TPH2-Cr R/Rosa26-tdTOMATO mice. (B-D) Repre entative electrophysiological trace (B), seline firing frequency (C), and resting memberane potential (D) of TPH2 (+) neurons in the DRN of male and female mice (n = 26 or 38). (E-L) The responses of firing frequency and resting membrane potential to EtOH treatment in the presence of vehicle or 500 μM 17β-Estradiol (n = 8 or 10). Mean ± SEM. (C-D) ***p<0.001, ****p<0.0001 in unpaired t-tests. (E-F and I-J) *p<0.05, **p<0.01, ***p<0.001, ****p<0.0001 in paired t-tests. (G-H and K-L) *p<0.05, ***p<0.001, ****p<0.0001 in one-way ANOVA analysis followed by post hoc Tukey tests.

### ERα agonist stimulates ERα^DRN^ neurons while ERβ agonist inhibits 5-HT^DRN^ neurons

To explore the intracellular mechanism for estrogenic action on 5-HT^DRN^ neurons, we recorded the responses of ERα^DRN^ or 5-HT^DRN^ neurons to treatment with the ERα agonist, propyl pyrazole triol (PPT), or the ERβ agonist, diarylpropionitrile (DPN), respectively. In both male and female mice, we found that PPT significantly increased firing frequency and depolarized resting membrane potential of ERα^DRN^ neurons (Fig. 5A-B), which is consistent with our previous report on the ERα-mediated stimulatory effects of PPT on 5-HT^DRN^ neurons^34^. This is also in line with the observations that most ERα-expressing neurons are 5-HT positive neurons (Fig. 2D). Conversely, DPN significantly decreased firing frequency and hyperpolarized resting membrane potential of 5-HT^DRN^ neurons (Fig. 5C-D), suggesting ERβ-mediated inhibition on 5-HT^DRN^ neurons^33^. Notably, around 70-77% of 5-HT^DRN^ neurons responded to DPN treatment (Fig. 5E), consistent with the observation that 70-73% of 5-HT^DRN^ neurons co-express ERβ (Fig. 2E). These results suggest antagonistic roles for ERα and ERβ expressed by 5-HT^DRN^ neurons in neural activity regulation.

**Fig. 5.**
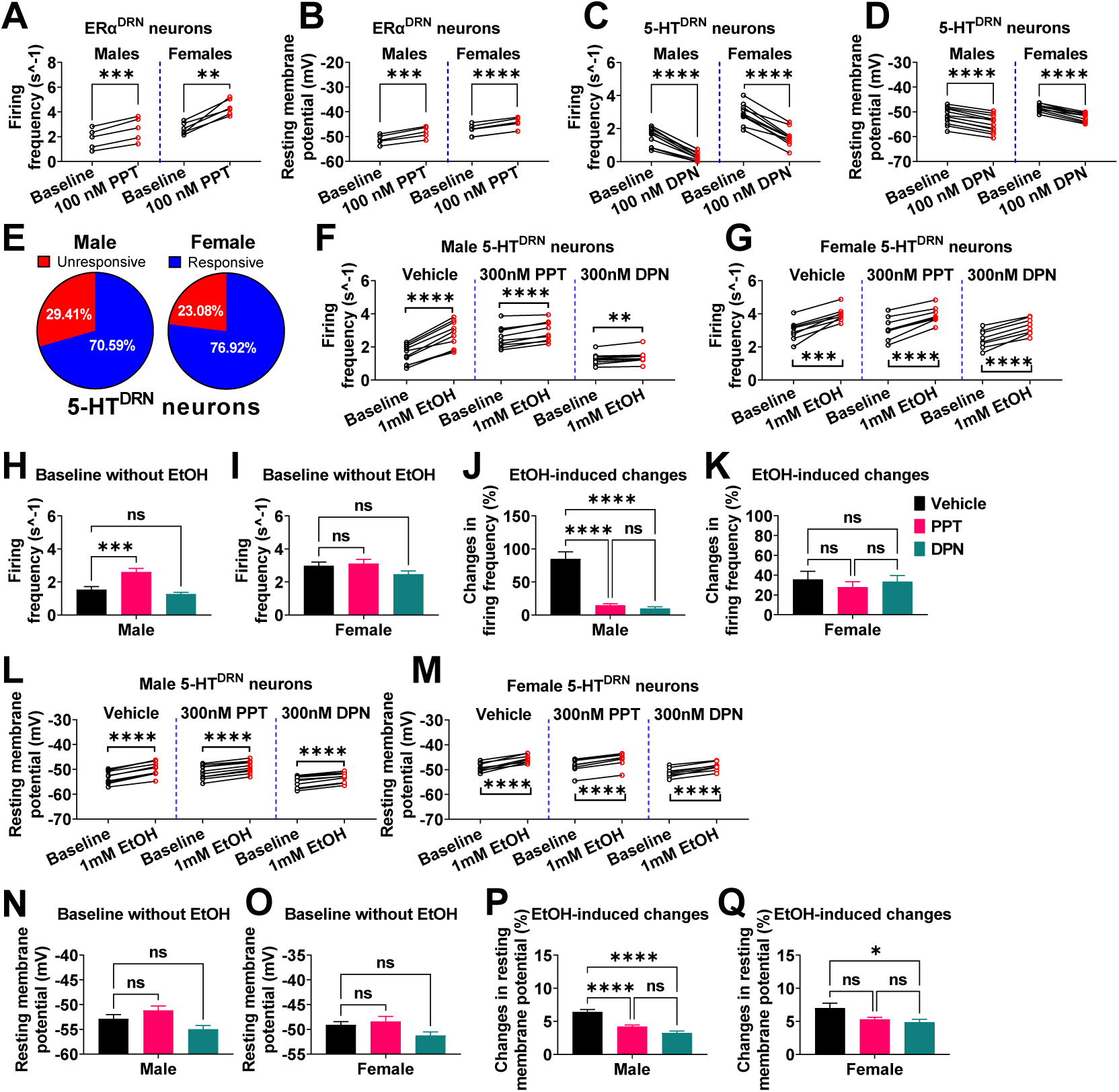
A selective agonist for ERα or ERβ attenuates EtOH-induced activation of 5-HTDRN neurons. (A-B) The firing frequency (A) and resting membrane potential (B) of ERα (+) neurons in the DRN of ERα-ZsGreen mice before and after a 1s puff of aCSF containing 100 nM PPT (n = 5 or 6). (C-E) The firing frequency (C), resting membrane potential (D), and responsive rate (E) of TPH2 (+) neurons in the DRN of TPH2-CreER/Rosa26-tdTOMATO mice before and after a 1s puff of aCSF containing 100 nM DPN (n = 10 or 11). (F-Q) The responses of firing frequency and resting membrane potential to EtOH treatment in the presence of vehicle, 300 nM PPT, or 300 nM DPN in TPH2 (+) neurons in the DRN of male and female TPH2-CreER/Rosa26-tdTOMATO mice (n = 8 or 10). Mean ± SEM. (A-B and D-E) **p<0.01, ***p<0.001, ****p<0.0001 in paired t-tests. (C and F) *p<0.05, **p<0.01, ***p<0.001, ****p<0.0001 in one-way ANOVA analysis followed by post hoc Tukey tests within each sex.

### Both ERα and ERβ agonists attenuate EtOH-induced excitation of 5-HT^DRN^ neurons

To identify the primary mediating receptor for E2’s inhibitory effects on ethanol-induced activation of 5-HT^DRN^ neurons, we pre-incubated DRN-containing brain slices from TPH2-CreER/Rosa26-tdTOMATO mice with either PPT or DPN. Subsequently, we tested the response of 5-HT^DRN^ neurons to ethanol treatment. We found that 1 mM ethanol puff significantly increased the firing frequency and depolarized resting membrane potential in 5-HT^DRN^ neurons regardless of sex or agonist treatment (Fig. 5F-G and L-M). At a baseline level without ethanol treatment, PPT bath incubation significantly increased the firing rate of 5-HT^DRN^ neurons in male but not in female mice (Fig. 5H-I), without changing resting membrane potential in both sexes (Fig. 5N-O). On the other hand, DPN failed to affect either firing frequency or resting membrane potential in both sexes (Fig. 5H-I and N-O). Notably, unlike puff treatment, in the current-clamp recording settings, 5-HT^DRN^ neurons were pre-incubated with PPT and DPN in the bath solutions, contributing to much more modest changes in baseline firing frequency and resting membrane potential. Importantly, the 5-HT^DRN^ neurons from males were unresponsive to ethanol in the presence of PPT or DPN (Fig. 5J and P). Conversely, in females, only DPN showed inhibitory effects on ethanol-induced changes in resting membrane potential (Fig. 5K and Q). These results suggest that both ERα and ERβ contribute to E2’s inhibitory effects on ethanol-induced activation of 5-HT^DRN^ neurons.

### Chemogenetic activation of ERα^DRN^ or ERβ^DRN^ neurons attenuates binge drinking in females

To directly test the regulatory effects of ERα^DRN^ neurons on binge drinking, we employed the DREADD method to selectively activate ERα^DRN^ neurons by stereotaxic injection of AAV-DIO-hM3Dq-mCherry into the DRN of *Esr1*-Cre mice (ERα-Dq^DRN^, Fig. 6A). To exclude the possible off-target effects of clozapine N-oxide (CNO) or its metabolites^35^, we included *Esr1*-Cre mice receiving the AAV-DIO-mCherry virus injections as controls (ERα-mCherry^DRN^). We observed specific expression of hM3Dq-mCherry in female ERα-Dq^DRN^ mice (Fig. 6B). Notably, the mCherry-positive neurons in the DRN were identified to be TPH neurons (Fig. 6C-E), and CNO treatment increased firing frequency and resting membrane potential of these neurons (Fig. 6F-J), validating the successful activation of ERα-expressing 5-HT^DRN^ neurons. We found that CNO-induced activation of ERα^DRN^ neurons reduced alcohol consumption in both 2-hour and 4-hour binge drinking sessions (Fig. 6K-L), suggesting an essential role of the ERα→5-HT pathway in binge drinking.

**Fig. 6.**
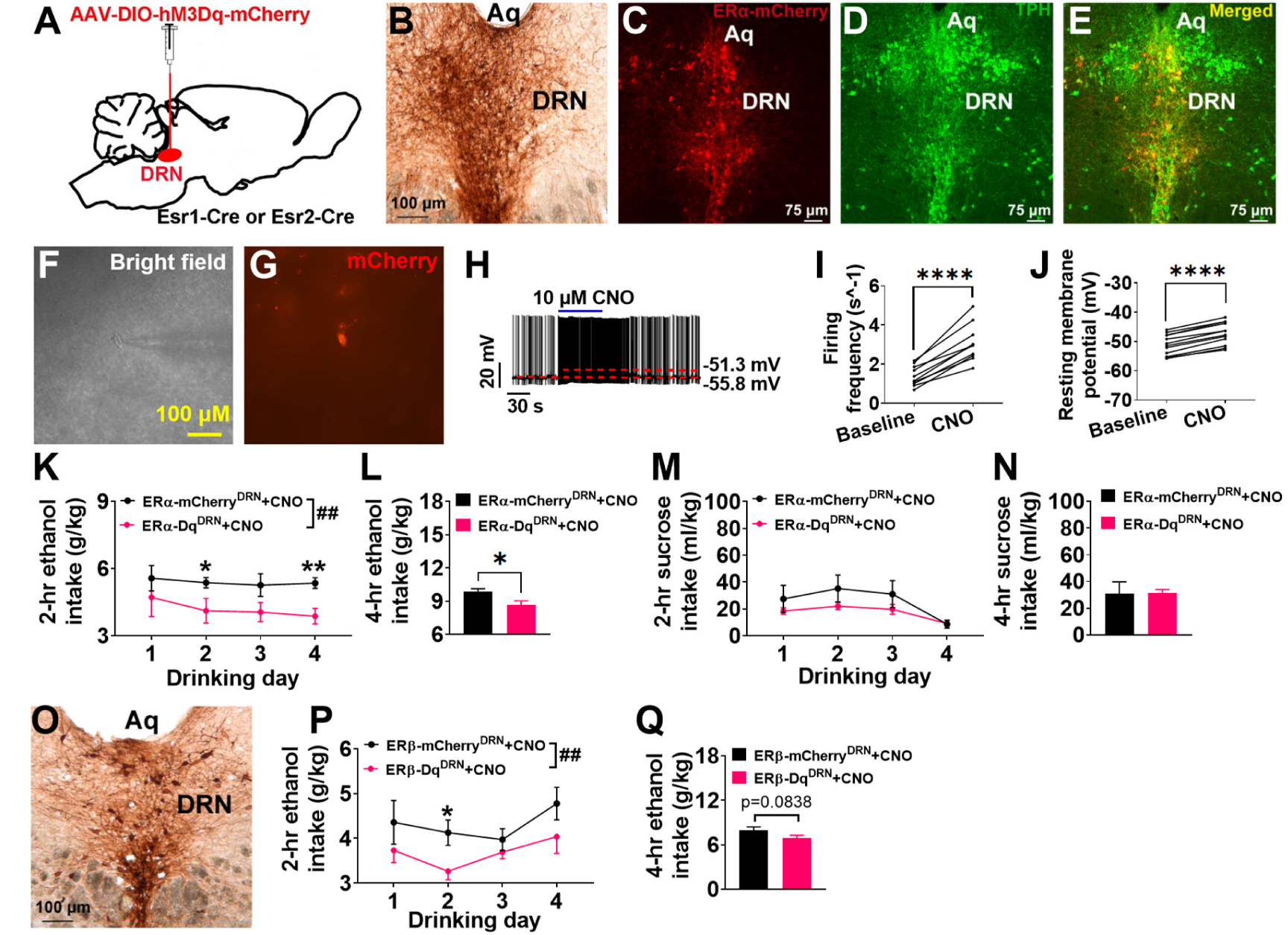
Activation of ERα^DRN^ or ERβ^DRN^ neurons decrease ethanol drinking. (A) Schematic of the experimental strategy using the AAV-DIO-hM3Dq-mCherry virus to selectively activate ERα^DRN^ or ERβ^DRN^ neurons in Esr1-Cre or Ers2-Cre female mice (ERα-Dq^DRN^ or ERβ-Dq^DRN^); Esr1-Cre or Esr2-Cre mice receiving the AAV-DIO-mCherry virus injections were used as controls (ERα-mCherry^DRN^ or ERβ-mCherry^DRN^). (B) Immunostaining of ERα-hM3Dq-mCherry in the DRN of female ERα-Dq^DRN^ mice. (C-E) Immunofluorescence staining of ERα-hM3Dq-mCherry (C), TPH (D), and merger (E) in the DRN of female ERα-Dq^DRN^ mice. (F-G) Micrographic images showing a recorded ERα-hM3Dq-mCherry (+) neurons in the DRN. (H-J) Representative traces and statistical analysis before and after CNO treatment. (K-L) Ethanol intake in g/kg over 2 hrs during 4 days of drinking (K) or during the final 4-hrs drinking session (L) after i.p. injection of CNO in female ERα-Dq^DRN^ or ERα-mCherry^DRN^ mice. (M-N) Sucrose intake in ml/kg over 2 hrs during 4 days of drinking (M) or during the final 4-hrs drinking session (N) after i.p. injection of CNO in female ERα-Dq^DRN^ or ERα-mCherry^DRN^ mice. (O) Immunostaining of ERβ-hM3Dq-mCherry in the DRN of female ERβ-Dq^DRN^ mice. (P-Q) Ethanol intake over 2 hrs during 4 days of drinking (P) or during the final 4-hrs drinking session (M) after i.p. injection of CNO in female ERβ-Dq^DRN^ or ERβ-mCherry^DRN^mice. Mean ± SEM. (E-F) ****p<0.0001 in paired t-tests. (G) ##P<0.01 in two way ANOVA analyses; *P<0.05, **P<0.01 in the following post hoc Bonferroni tests. (H) *P<0.05 in non-paired T-tests.

To determine whether the effects of activation of ERα^DRN^ neurons on binge-like drinking were specific to ethanol or might extend to other rewarding substances, we performed the DID using 2% sucrose (instead of ethanol). There were no significant effects of ERα^DRN^ activation for sucrose consumption during the 2-hour sessions or during the final 4-hour session (Fig. 6M-N), demonstrating that activation of ERα^DRN^ neurons in female mice does not affect sucrose-induced reward effects. Finally, to determine whether the inhibitory effect of ERα^DRN^ activation on alcohol consumption is ERα-specific, we injected AAV-DIO-hM3Dq-mCherry into the DRN of Esr2-Cre mice (Fig. 6O) and tested the effects of ERβ^DRN^ activation on the ethanol drinking in the DID. We found that activation of ERβ^DRN^ neurons in female mice also significantly decreased alcohol consumption during the 2-hour sessions and tended to reduce ethanol drinking during the final 4-hour session (Fig. 6P-O). These results suggest a regulatory role for ERα- and ERβ-expression neurons in the DRN in ethanol binge drinking.

## Discussion

Our findings demonstrate a regulatory role of estrogen/ERs/5-HT^DRN^ signaling in binge drinking. To our knowledge, this is the first report regarding the role of estrogenic serotonin system in controlling alcohol binge drinking. The essential function of this signaling is first supported by the sex-specific alterations in mRNA expression of ERα and 5-HT-related genes in the DRN induced by chronic alcohol DID tests. We further provided *ex vivo* patch-clamp evidence that ethanol sex dimorphically depolarizes 5-HT^DRN^ neurons and that pharmacological activation of ERα or ERβ attenuates ethanol-induced activation of 5-HT neurons. Finally, we showed that DREADD stimulation of ERα^DRN^ or ERβ^DRN^ neurons reduces binge-like ethanol drinking. These findings implicate a potential role of estrogen signaling in 5-HT^DRN^ neurons in sex dimorphism in binge-like alcohol drinking.

Although many studies have shown an association between estrogen signaling and the brain’s 5-HT system, a key modulator of alcohol intake^23-25^, the potential regulatory role of estrogenic 5-HT signaling in alcohol drinking has not been thoroughly studied. Anatomically, ERα and ERβ are expressed in the 5-HT^DRN^ neurons of male and female mice^30^. We further quantified the expression pattern of ERα, ERβ, and TPH in the DRN and found most 5-HT^DRN^ neurons are positive for ERα or ERβ. Notably, a large portion of the 5-HT^DRN^ neurons co-express both ERα and ERβ. This drove us to separately test the electrophysiological function of ERα or ERβ expressed by 5-HT^DRN^ neurons. In line with our previous reports^32,33^, we found that estrogens stimulate 5-HT^DRN^ neurons through ERα while inhibiting 5-HT^DRN^ neurons through ERβ, suggesting antagonistic effects of ERα and ERβ expressed by 5-HT^DRN^ neurons. It is puzzling that two receptors with opposite regulatory effects on neural activity are expressed in the same neurons. Interestingly, several previous studies have consistently observed that neurons in different brain regions, including the medial amygdala (MeA), the bed nucleus of the stria terminalis (BNST), and the preoptic area of the hypothalamus (POAH), co-express both ERα and ERβ^36,37^. It has been proposed that the presence of both ERs may allow the neurons to accommodate different physiological/pathological conditions by changing ERα/ERβ ratio and responding differentially to the actions of estrogen^37^.

Consistent with this point of view, in male mice, expression of *Esr1* was significantly increased by chronic alcohol DID, resulting in a much lower *Esr2*/*Esr1* ratio in the DRN. This alcohol-induced lower ratio was associated with higher 5-HT synthesis and reuptake, as indicated by higher mRNA expression of *Sert* and *Pet1* in the DRN of males induced by chronic alcohol DID. These findings suggest that during the transition from alcohol-naive to chronic alcohol DID condition, estrogens/ERα-mediated activation may override estrogens/ERβ-mediated inhibition in 5-HT^DRN^ neurons, contributing to the higher 5-HT transmission induced by chronic alcohol DID in males.

Notably, aromatase, the key enzyme for estrogen synthesis, is highly expressed in several male brain regions, including MeA, BNST, and POAH^38^. These brain regions could project and release estrogen in the DRN. Aromatase can transform testosterone into E2, which means that even in the male brain, the DRN could be exposed to high levels of E2 produced locally from testosterone. Consistently, it has been shown that 5-HT secretion from 5-HT^DRN^ neurons in males is remarkably decreased by aromatase inhibition^39^. Male estrogen signaling may play an essential role in the serotonergic adaptation induced by chronic alcohol DID in males.

Another interesting observation is the sex difference in the *Esr2*/*Esr1* ratio in the DRN of alcohol-naïve mice. Compared to males, female naïve mice have a much lower *Esr2*/*Esr1* ratio and higher mRNA expression of key enzymes for 5-HT synthesis and reuptake in the DRN. Consistently, we also observe sex differences in the intrinsic electrophysiological properties of 5-HT^DRN^ neurons in the baseline condition without alcohol exposure. Female 5-HT^DRN^ neurons showed much higher firing rate and depolarized resting membrane potential, indicating a higher baseline activity in female 5-HT^DRN^ neurons compared to that in males.

After chronic alcohol DID, the difference in *Esr2*/*Esr1* ratio between males and females was diminished, associated with abolished sex differences in mRNA expression of *Sert, Pet1*, and *Tph2*. These findings support that the dynamic counterbalance of ERα and ERβ in the 5-HT^DRN^ neurons may play a physiological role in the regulation of serotonergic tone in the DRN during chronic alcohol DID exposure.

Notably, acute binge drinking causes a burst release of central 5-HT, and increased brain 5-HT inhibits alcohol consumption in humans and rodents^23-26^. These findings suggest that the alcohol-induced activation of 5-HT neurons may be an essential component of a negative feedback loop to decrease acute alcohol consumption. We speculate that the rapid responses of 5-HT^DRN^ neurons to alcohol are distinct in males and females. These differences may contribute to the sex dimorphism in the alcohol binge-like drinking behavior. To test this hypothesis, we compared the acute electrophysiological response of male and female ERα^DRN^ neurons to ethanol treatment. Consistent with our previous findings^34^, we confirm that most ERα-expressing cells within the DRN are 5-HT neurons. Like 5-HT^DRN^ neurons, female ERα^DRN^ neurons also showed high baseline neuronal firing activity dynamics than male ERα^DRN^ neurons. Importantly, we found that male 5-HT^DRN^ neurons showed a lower threshold responding dose (0.5 mM in males vs. 1 mM in females) and higher percentage changes in firing frequency and resting membrane potential when incubated with the same amount of ethanol. These results suggest that male ERα^DRN^ neurons are more sensitive to ethanol treatment compared to female ERα^DRN^ neurons. We postulate that lower baseline neuronal firing activity of male ERα^DRN^ neurons leads to higher burse increases of 5-HT release, contributing to earlier ethanol drinking termination.

Sex differences in the 5-HT system and the regulatory effects of estrogens on 5-HT neuron activity have been demonstrated for decades^31,40-42^. It has been shown that estrogen-mediated sex differences in the 5-HT system contribute to the greater susceptibility of women to many affective behavior disorders, such as premenstrual syndrome, postpartum depression, and postmenopausal depression^31,41^. We speculate that estrogen signaling may partially mediate the sex difference in ethanol-induced excitation of 5-HT^DRN^ neurons. In supporting this view, we observed that E2 treatment significantly attenuated the ethanol-induced excitation of 5-HT^DRN^ neurons in males but not in females, suggesting an estrogen-mediated attenuation of ethanol-induced 5-HT burst release. We further demonstrated that both ERα and ERβ agonists consistently produced similar inhibitory effects on ethanol-induced firing activity changes of 5-HT^DRN^ neurons, despite antagonistic effects on activity dynamics of 5-HT^DRN^ neurons. These results suggest distinct mechanisms for ERs’ effects on baseline activity and ethanol-induced burst firing. These findings support the notion that estrogens acutely increase alcohol binge drinking by reducing the responsiveness of 5-HT neurons to ethanol.

Another critical observation supporting our models is that chemogenetic activation of ERα-expressing 5-HT^DRN^ neurons in females resulted in a significant decrease in alcohol consumption without affecting sucrose intake in the DID tests. Notably, activation of ERβ^DRN^ neurons also induced similar inhibition on ethanol intake, consistent with the observation that a subpopulation of 5-HT^DRN^ neurons co-expressing ERα and ERβ. These findings indicate that alcohol-induced acute stimulation of 5-HT^DRN^ neurons may serve as a defense mechanism to limit alcohol drinking. Estrogens-mediated sex differences in excitation of 5-HT^DRN^ neurons induced by ethanol may partially contribute to the sex dimorphism in binge-like drinking behavior in mice.

In conclusion, we demonstrated that the DRN 5-HT system differs significantly between sexes and undergoes profound sex-specific changes during chronic alcohol DID. Furthermore, estrogens may inhibit ethanol-induced acute excitation of 5-HT^DRN^ neurons, resulting in higher levels of binge drinking in females. Notably, different neuroactive mechanisms seem to be involved in the regulatory effects of estrogens on baseline activity and ethanol-induced burst firing. Additional studies are warranted to know what afferent or intracellular channels modulate the responsiveness of 5-HT^DRN^ neurons to ethanol. Whatever the exact mechanism, the significantly lower responses of female 5-HT^DRN^ neurons to ethanol may reduce a burst of 5-HT-mediated neurotransmission during alcohol binge drinking, thus diminishing a protective mechanism for alcohol overconsumption. In particular, it could partially explain the higher binge-like drinking behavior in female mice and sex differences in human alcohol drinking behavior.

## Materials and methods

### Animals

Several transgenic mouse lines were maintained on a C57BL/6J background. These lines include *Esr1*-Cre mice (#017911, Jackson Laboratory, Bar Harbor, ME), *Esr2*-Cre mice (#30158, Jackson Laboratory), TPH2-iCreER (#016584, Jackson Laboratory), Rosa26-LSL-tdTOMATO (#007914, Jackson Laboratory), ERβ-EGFP (#030078-UCD, MMRRC at UC Davis), and ERα-ZsGreen^43^. C57BL/6J male and female mice were purchased from The Jackson Laboratory. TPH2-iCreER and Rosa26-LSL-tdTOMATO were crossed to generate TPH2-iCreER/Rosa26-LSL-tdTOMATO for electrophysiological recording. *Esr2*-Cre and Rosa26-LSL-tdTOMATO were crossed to generate *Esr2*-Cre/Rosa26-LSL-tdTOMATO for immunofluorescent staining. Mice were housed in a temperature-controlled environment at 22 °C-24 °C on a 12-hour light/dark cycle (light off at 6 pm) or a 12-hour reversed light/dark cycle (light off at 10 am). Unless otherwise stated, the mice were fed ad libitum with standard mouse chow (6.5 % fat, #2920, Harlan-Teklad, Madison, WI) and water. Care of all animals and procedures were approved by Pennington Biomedical Research Center (PBRC) and The University of Illinois at Chicago Institutional Animal Care and Use Committees.

### Chronic Drinking in the dark (DID)

Both male and female C57BL/6J mice were subjected to 9-week long water or ethanol DID. The 4-day DID test was performed as described previously with minor modification^44^. Briefly, mice were individually housed in a 12-hour reversed light/dark cycle room (lights off at 10 AM and on at 10 PM) for two weeks before behavioral testing. For water DID group, water consumption was measured by replacing the water bottle 3 hours into the dark cycle with a single-sipper tube containing only water. On the first three days (Monday, Tuesday, and Wednesday), mice were given access to the sipper tube for 2 hours. On the fourth day, mice were given access to the sipper tube for 4 hours. Total consumption of water over the 2-hour (1^st^-4^th^ days) and 4-hour (4^th^ day) period was measured in each individual. Similarly, for ethanol DID group, ethanol consumption was measured by the ethanol DID test. Mice underwent a DID test identical to the water consumption test, except the sipper tube contained 20% ethanol in water instead of only water. The water or ethanol DID was performed on Mon-Thurs each week for 9 weeks, with no ethanol access on Fri-Sun. Sixteen hours after the last drinking session (9 AM), mice were euthanized via cardiac puncture under anesthesia. The dorsal raphe nucleus (DRN) was punched out and stored in -80°C for further analysis.

### DREADD stimulation of ERα^DRN^ or ERβ^DRN^ neuron

Female *Esr1*-Cre or *Esr2*-Cre were stereotaxically injected with 400 nL AAV-DIO-hM3Dq-mCherry (#GVVC-AAV-130, Stanford Virus Core) or AAV-DIO-mCherry (UNC Vector Core) into the DRN (4.65 mm posterior, 0 mm lateral and 3.60 mm ventral to the Bregma, based on Franklin & Paxinos Mouse Brain Atlas) at eight weeks of age. Four weeks after the surgery, all mice were singly housed and acclimated into a 12-hour reversed light/dark cycle room. Two weeks after, ethanol DID was performed as described above. Clozapine N-oxide (CNO, 3 mg/kg) was intraperitoneal (i.p.) injected 2 hours into the dark cycle in each drinking session day. One week later, sucrose consumption was measured by the sucrose DID test. All the procedures were identical to the ethanol DID test, except the sipper tube contained 2 percent sucrose in water. After studies, all mice were perfused with 10% formalin, and the brains were collected and sliced. Sections were collected for immunohistochemistry of mCherry. Briefly, brain sections were incubated with rabbit anti-DsRed antibody (1:1,000; #632496, Takara Bio., Mountain View, CA) at room temperature overnight, followed by the biotinylated donkey anti-rabbit secondary antibody (1:1000, #711-067-003, Jackson ImmunoResearch, West Grove, PA) for 1 hour. Sections were then incubated in the avidin-biotin complex (1:1000, PK-6100, Vector Laboratories) and followed by 0.04% 3, 3′-diaminobenzidine in 0.01% hydrogen peroxide. After dehydration through graded ethanol, the slides were immersed in xylene and cover slipped. Bright-field images were analyzed.

Another aliquot of sections from *Esr1*-Cre mice injected with AAV-DIO-hM3Dq-mCherry were used for immunofluorescent staining of TPH. Briefly, brain sections were incubated with sheep anti-TPH antibody (1:1000, #AB1541, Millipore) at room temperature overnight, followed by the Alexa Fluor 488-conjugated donkey anti-sheep (1:500; #713-545-003, Jackson ImmunoResearch) for 2 hours. Fluorescent images were obtained using a Leica DM5500 fluorescence microscope with OptiGrid structured illumination configuration.

### Real-time RT-PCR in DRN

Total mRNAs were extracted using TRIzol (#15596018, Invitrogen, Carlsbad, CA). SYBR Green quantitative PCR (qPCR) was performed, as described previously^45,46^. Primer sequences were listed in Table 1. Results were normalized by the expression of *Gapdh* as the reference gene.

**Table 1.**
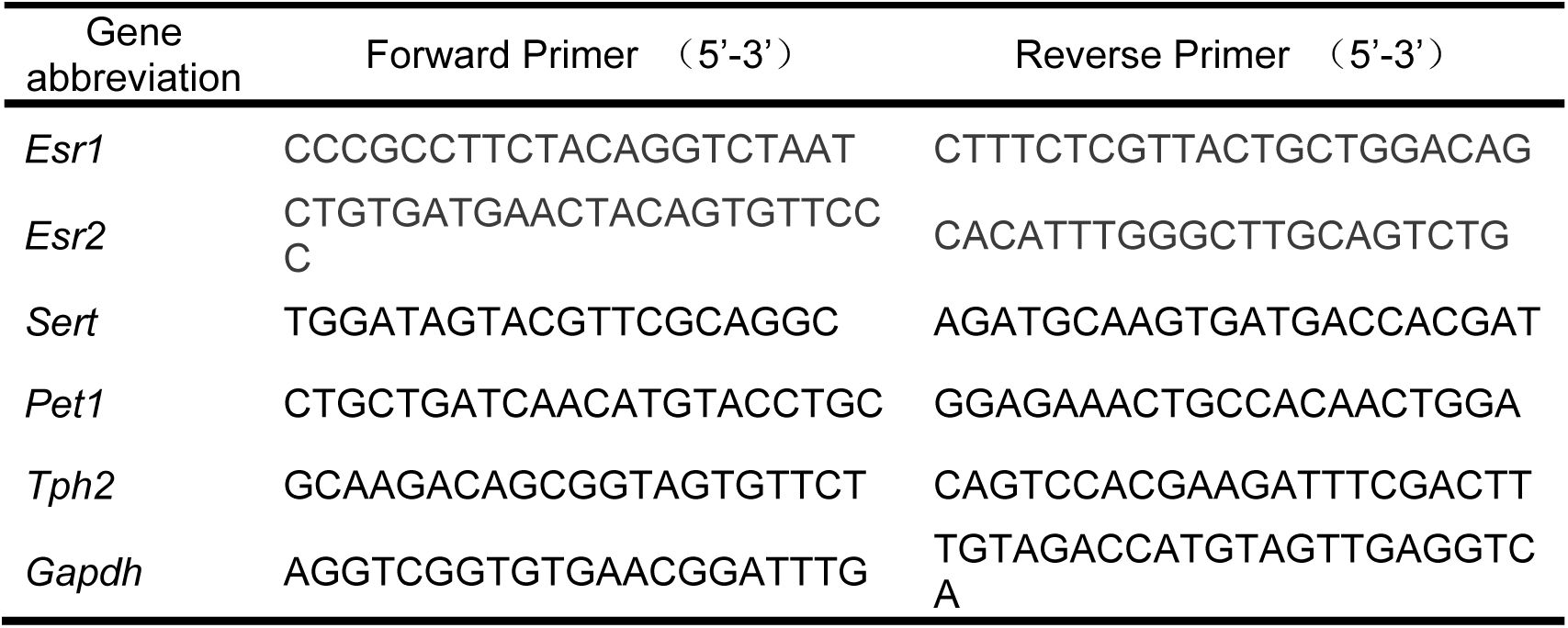
qPCR primer sequences of related genes

### Co-staining of ERα, ERβ, and TPH in the DRN

Both male and female *Esr2*-Cre/Rosa26-LSL-tdTOMATO mice were perfused with 10% formalin at 8 weeks of age. Female mice were perfused at diestrus when circulating estrogens were low^47^. The brains were collected and sliced. Sections were collected for double staining of ERα and TPH. Briefly, brain sections were incubated with rabbit anti-ERα antibody (1:5,000; #06-935, Millipore, Burlington, MA) at room temperature overnight, followed by the Alexa Fluor 647-conjugated donkey anti-rabbit secondary antibody (1:500, #711-605-152, Jackson ImmunoResearch) for 2 hours. Sections were then incubated in sheep anti-TPH antibody (1:1000, #AB1541, Millipore) overnight, followed by the Alexa Fluor 488-conjugated donkey anti-sheep (1:500; #713-545-003, Jackson ImmunoResearch) for 2 hours. Fluorescent images were obtained using a Leica DM5500 fluorescence microscope with OptiGrid structured illumination configuration.

Similarly, double immunofluorescent staining of ERα and TPH was performed in DRN sections from a female ERβ-EGFP mouse. All staining procedures are identical except using the Alexa Fluor 594-conjugated donkey anti-sheep (1:500; #713-585-003, Jackson ImmunoResearch) as the secondary antibody for TPH.

### Electrophysiology

The electrophysiological responses of identified ERα neurons in the DRN to ethanol treatment were investigated in ERα-ZsGreen mice as previously described^34^. Briefly, whole-cell patch-clamp recordings were performed on identified green fluorescent neurons in the brain slices containing DRN from ERα-ZsGreen mice. Six to twelve-week-old mice were deeply anesthetized with isoflurane and transcardially perfused with an ice-cold, carbogen-saturated (95% O2, 5% CO2) sucrose-based cutting solution (pH 7.3), containing 10 mM NaCl, 25 mM NaHCO3, 195 mM Sucrose, 5 mM Glucose, 2.5 mM KCl, 1.25 mM NaH2PO4, 2 mM Na pyruvate, 0.5 mM CaCl2, 7 mM MgCl2. The entire brain was removed and coronally cut into slices (250 μm) with a Microm HM 650V vibratome (Thermo Fisher Scientific, Waltham, MA). Then, the DRN-containing slices were incubated in oxygenated aCSF (adjusted to pH7.3) containing (in mM) 126 NaCl, 2.5 KCl, 2.4 CaCl2, 1.2 NaH2PO4, 1.2 MgCl2, 11.1 glucose, and 21.4 NaHCO3 for one hour at 34°C.

Slices were transferred to the recording chamber and perfused at 34ºC in oxygenated aCSF at a flow rate of 1.8-2 mL/min. ZsGreen-labeled ERα^DRN^ neurons were visualized using epifluorescence and IR-DIC imaging. The intracellular solution (adjusted to pH 7.3) contained the following (in mM): 128 K gluconate, 10 KCl, 10 HEPES, 0.1 EGTA, 2 MgCl2, 0.05 Na-GTP and 0.05 Mg-ATP. Recordings were made using a MultiClamp 700B amplifier (Molecular Devices, Sunnyvale, CA, United States), sampled using Digidata 1440A, and analyzed offline with pClamp 10.3 software (Molecular Devices). Series resistance was monitored during the recording, and the values were generally < 10 MΩ and were not compensated. The liquid junction potential (LJP) was +12.5 mV and was corrected after the experiment. Data was excluded if the series resistance increased dramatically during the experiment or without overshoot for the action potential. Currents were amplified, filtered at 1 kHz, and digitized at 20 kHz. The current clamp was engaged in testing neuronal firing and resting membrane potential before and after a 1s puff of aCSF containing vehicle or ethanol (0.1, 0.5, 1, 10, or 50 mM).

To study the effect of a selective ERβ agonist, diarylpropionitrile (DPN), on the activity of 5-HT neurons, a cohort of both male and female TPH2-iCreER/Rosa26-LSL-tdTOMATO mice were generated. Tamoxifen (0.2 mg/g body weight) was i.p. injected to induce expression of tdTOMATO in 5-HT neurons four weeks before the brain slice recording. The current clamp was engaged in testing neural firing and resting membrane potential of tdTOMATO-labeled 5-HT neurons in the DRN before and after a 1s puff of aCSF containing a vehicle or 100 nM DPN.

In a different cohort of TPH2-iCreER/Rosa26-LSL-tdTOMATO mice, the brain slices were pre-incubated with aCSF containing vehicle, 17β-estradiol (E2, 500uM), propyl pyrazole triol (PPT, 300nM), or DPN (300nM). PPT is a selective ERα agonist, while DPN is a selective ERβ agonist. The current clamp was engaged in testing neural firing and resting membrane potential before and after a 1s puff of aCSF containing a vehicle or 1 mM ethanol in the presence of vehicle, E2, PPT, or DPN.

To assess the effect of CNO on ERα neurons, 8-to 10-week-old Esr1-Cre mice were injected with AAV-DIO-hM3Dq-mCherry into the DRN 2-3 weeks before recording. Brain slices were prepared and CNO was applied to the bath solution through perfusion as previously described^48^. Effects of CNO (10 μM) on membrane potential and firing frequency of mCherry-labeled ERα in the DRNA were electrophysiologically recorded.

### Statistics

Statistical analyses were performed using GraphPad Prism 7.0 statistics software (San Diego, CA USA). Statistical analyses methods were chosen based on the design of each experiment and indicated in the figure legends. The data were presented as mean ± SEM. P ≤0.05 was considered statistically significant.

## Author Contributions

VT and BF are the main contributors in the conduct of the study, data collection and analysis, data interpretation, and manuscript writing. XY, PN, SS, LI, HY, PL, LC, PL, MK, and DD contributed to the conduct of the study. CW, AWL, YH, and PX contributed to the study design, data interpretation, and manuscript writing.

## Acknowledgments

This work was supported by grants from NIH (R01 DK123098, P30 DK020595 to PX; P20 GM135002, R01 DK129548 to YH; T32 AA026577 to VT; P50 AA022538, U01 AA020912, and R01 AA027231 to AWL; K01 DK119471 to CW), USDA/CRIS (3092-51000-062-04(B)S to CW), AHA (915789 to VT; 19CDA34660335 to CW; 20POST35120600 to YH), and DOD (Innovative Grant W81XWH-20-1-0075 to PX). The authors wish to thank the Laboratory of Animal Center of Pennington Biomedical Research Center at Louisiana State University, and The University of Illinois at Chicago for invaluable help in mouse colony maintenance.

## References

1. Kanny, D., Naimi, T.S., Liu, Y., Lu, H. & Brewer, R.D. Annual Total Binge Drinks Consumed by U.S. Adults, 2015. Am J Prev Med 54, 486–496 (2018).

2. Esser, M.B., et al. Prevalence of alcohol dependence among US adult drinkers, 2009-2011. Prev Chronic Dis 11, E206 (2014).

3. Kanny, D., et al. Binge drinking - United States, 2011. MMWR Suppl 62, 77–80 (2013).

4. White, A., et al. Converging Patterns of Alcohol Use and Related Outcomes Among Females and Males in the United States, 2002 to 2012. Alcohol Clin Exp Res 39, 1712–1726 (2015).

5. O’Keefe, J.H., Bybee, K.A. & Lavie, C.J. Alcohol and cardiovascular health: the razor-sharp double-edged sword. J Am Coll Cardiol 50, 1009–1014 (2007).

6. Zakhari, S. & Li, T.K. Determinants of alcohol use and abuse: Impact of quantity and frequency patterns on liver disease. Hepatology 46, 2032–2039 (2007).

7. Wilsnack, S.C., Wilsnack, R.W. & Kantor, L.W. Focus on: women and the costs of alcohol use. Alcohol Res 35, 219–228 (2013).

8. Petit, G., Maurage, P., Kornreich, C., Verbanck, P. & Campanella, S. Binge drinking in adolescents: a review of neurophysiological and neuroimaging research. Alcohol Alcohol 49, 198–206 (2014).

9. Wilhelm, C.J., et al. Astrocyte Dysfunction Induced by Alcohol in Females but Not Males. Brain Pathol 26, 433–451 (2016).

10. Muti, P., et al. Alcohol consumption and total estradiol in premenopausal women. Cancer Epidemiol Biomarkers Prev 7, 189–193 (1998).

11. Frydenberg, H., et al. Alcohol consumption, endogenous estrogen and mammographic density among premenopausal women. Breast Cancer Res 17, 103 (2015).

12. Hartman, T.J., et al. Alcohol Consumption and Urinary Estrogens and Estrogen Metabolites in Premenopausal Women. Horm Cancer 7, 65–74 (2016).

13. Martin, C.A., Mainous, A.G., 3rd, Curry, T. & Martin, D. Alcohol use in adolescent females: correlates with estradiol and testosterone. Am J Addict 8, 9–14 (1999).

14. Reid, M.L., Hubbell, C.L. & Reid, L.D. A pharmacological dose of estradiol can enhance appetites for alcoholic beverages. Pharmacol Biochem Behav 74, 381–388 (2003).

15. Mackie, A.R., et al. Alcohol consumption negates estrogen-mediated myocardial repair in ovariectomized mice by inhibiting endothelial progenitor cell mobilization and function. J Biol Chem 288, 18022–18034 (2013).

16. Satta, R., Hilderbrand, E.R. & Lasek, A.W. Ovarian Hormones Contribute to High Levels of Binge-Like Drinking by Female Mice. Alcohol Clin Exp Res 42, 286–294 (2018).

17. Becker, H.C., Anton, R.F., De Trana, C. & Randall, C.L. Sensitivity to ethanol in female mice: effects of ovariectomy and strain. Life Sci 37, 1293–1300 (1985).

18. Ford, M.M., Eldridge, J.C. & Samson, H.H. Ethanol consumption in the female Long-Evans rat: a modulatory role of estradiol. Alcohol 26, 103–113 (2002).

19. Mello, N.K., Bree, M.P. & Mendelson, J.H. Alcohol and food self-administration by female Macaque monkeys as a function of menstrual cycle phase. Physiol Behav 36, 959–966 (1986).

20. Vandegrift, B.J., et al. Estrogen Receptor alpha Regulates Ethanol Excitation of Ventral Tegmental Area Neurons and Binge Drinking in Female Mice. J Neurosci 40, 5196–5207 (2020).

21. Vandegrift, B.J., You, C., Satta, R., Brodie, M.S. & Lasek, A.W. Estradiol increases the sensitivity of ventral tegmental area dopamine neurons to dopamine and ethanol. PLoS One 12, e0187698 (2017).

22. You, C., Vandegrift, B. & Brodie, M.S. Ethanol actions on the ventral tegmental area: novel potential targets on reward pathway neurons. Psychopharmacology (Berl) 235, 1711–1726 (2018).

23. LeMarquand, D., Pihl, R.O. & Benkelfat, C. Serotonin and alcohol intake, abuse, and dependence: clinical evidence. Biol Psychiatry 36, 326–337 (1994).

24. LeMarquand, D., Pihl, R.O. & Benkelfat, C. Serotonin and alcohol intake, abuse, and dependence: findings of animal studies. Biol Psychiatry 36, 395–421 (1994).

25. Pettinati, H.M. Use of serotonin selective pharmacotherapy in the treatment of alcohol dependence. Alcohol Clin Exp Res 20, 23A–29A (1996).

26. Litten, R.Z., Allen, J. & Fertig, J. Pharmacotherapies for alcohol problems: a review of research with focus on developments since 1991. Alcohol Clin Exp Res 20, 859–876 (1996).

27. Gundlah, C., et al. Estrogen receptor-beta regulates tryptophan hydroxylase-1 expression in the murine midbrain raphe. Biol Psychiatry 57, 938–942 (2005).

28. Osterlund, M.K. Underlying mechanisms mediating the antidepressant effects of estrogens. Biochim Biophys Acta 1800, 1136–1144 (2010).

29. Donner, N. & Handa, R.J. Estrogen receptor beta regulates the expression of tryptophan-hydroxylase 2 mRNA within serotonergic neurons of the rat dorsal raphe nuclei. Neuroscience 163, 705–718 (2009).

30. Mitra, S.W., et al. Immunolocalization of estrogen receptor beta in the mouse brain: comparison with estrogen receptor alpha. Endocrinology 144, 2055–2067 (2003).

31. Robichaud, M. & Debonnel, G. Oestrogen and testosterone modulate the firing activity of dorsal raphe nucleus serotonergic neurones in both male and female rats. J Neuroendocrinol 17, 179–185 (2005).

32. Cao, X., et al. Estrogens stimulate serotonin neurons to inhibit binge-like eating in mice. J Clin Invest 124, 4351–4362 (2014).

33. Saito, K., Cao, X., He, Y. & Xu, Y. Progress in the molecular understanding of central regulation of body weight by estrogens. Obesity (Silver Spring) 23, 919–926 (2015).

34. Cao, X., et al. Estrogens stimulate serotonin neurons to inhibit binge-like eating in mice. J Clin Invest 124, 4351–4362 (2014).

35. Gomez, J.L., et al. Chemogenetics revealed: DREADD occupancy and activation via converted clozapine. Science 357, 503–507 (2017).

36. Shughrue, P.J., Lane, M.V. & Merchenthaler, I. Comparative distribution of estrogen receptor-alpha and -beta mRNA in the rat central nervous system. J Comp Neurol 388, 507–525 (1997).

37. Shughrue, P.J., Scrimo, P.J. & Merchenthaler, I. Evidence for the colocalization of estrogen receptor-beta mRNA and estrogen receptor-alpha immunoreactivity in neurons of the rat forebrain. Endocrinology 139, 5267–5270 (1998).

38. Wu, M.V., et al. Estrogen masculinizes neural pathways and sex-specific behaviors. Cell 139, 61–72 (2009).

39. Bethea, C.L., Reddy, A.P., Robertson, N. & Coleman, K. Effects of aromatase inhibition and androgen activity on serotonin and behavior in male macaques. Behav Neurosci 127, 400–414 (2013).

40. Klink, R., Robichaud, M. & Debonnel, G. Gender and gonadal status modulation of dorsal raphe nucleus serotonergic neurons. Part I: effects of gender and pregnancy. Neuropharmacology 43, 1119–1128 (2002).

41. Dominguez, R., Cruz-Morales, S.E., Carvalho, M.C., Xavier, M. & Brandao, M.L. Sex differences in serotonergic activity in dorsal and median raphe nucleus. Physiol Behav 80, 203–210 (2003).

42. Nishizawa, S., et al. Differences between males and females in rates of serotonin synthesis in human brain. Proc Natl Acad Sci U S A 94, 5308–5313 (1997).

43. Saito, K., et al. Visualizing estrogen receptor-alpha-expressing neurons using a new ERalpha-ZsGreen reporter mouse line. Metabolism 65, 522–532 (2016).

44. Rhodes, J.S., Best, K., Belknap, J.K., Finn, D.A. & Crabbe, J.C. Evaluation of a simple model of ethanol drinking to intoxication in C57BL/6J mice. Physiol Behav 84, 53–63 (2005).

45. Xu, Y., et al. Distinct hypothalamic neurons mediate estrogenic effects on energy homeostasis and reproduction. Cell Metab 14, 453–465 (2011).

46. Wang, C., et al. TAp63 contributes to sexual dimorphism in POMC neuron functions and energy homeostasis. Nat Commun 9, 1544 (2018).

47. Hong, K. & Choi, Y. Role of estrogen and RAS signaling in repeated implantation failure. BMB Rep 51, 225–229 (2018).

48. Zhu, C., et al. Heparin Increases Food Intake through AgRP Neurons. Cell Rep 20, 2455–2467 (2017).

